# Molecular data from Orthonectid worms show they are highly degenerate members of phylum Annelida not phylum Mesozoa

**DOI:** 10.1101/235549

**Authors:** Philipp H. Schiffer, Helen E. Robertson, Maximilian J. Telford

## Abstract

The Mesozoa are a group of tiny, extremely simple, vermiform endoparasites of various marine animals (Fig. 1). There are two recognised groups within the Mesozoa: the Orthonectida (Fig. 1a,b; with a few hundred cells including a nervous system made up of just 10 cells [1]) and the Dicyemids (Fig. 1c; with at most 42 cells [2]). They are classic ‘Problematica’ [3] - the name Mesozoa suggests an evolutionary position intermediate between Protozoa and Metazoa (animals) [4] and implies their simplicity is a primitive state, but molecular data have shown they are members of Lophotrochozoa within Bilateria [5-8] which would mean they derive from a more complex ancestor. Their precise phylogenetic affinities remain uncertain, however, and ascertaining this is complicated by the very fast evolution observed in genes from both groups, leading to the common systematic error of Long Branch Attraction (LBA) [9]. Here we use mitochondrial and nuclear gene sequence data, and show beyond doubt that both dicyemids and orthonectids are members of the Lophotrochozoa. Carefully addressing the effects of systematic errors due to unequal rates of evolution, we show that the phylum Mesozoa is polyphyletic. While the precise position of dicyemids remains unresolved within Lophotrochozoa, we unequivocally identify orthonectids as members of the phylum Annelida. This result reveals one of the most extreme cases of body plan simplification in the animal kingdom; our finding makes sense of an annelid-like cuticle in orthonectids [1] and suggests the circular muscle cells repeated along their body [10] may be segmental in origin.

## Results

Using a new assembly of available genomic and transcriptomic sequence data we identified an almost complete mitochondrial genome from *Intoshia linei* (2 ribosomal RNAs, 20 transfer RNAs and all protein coding genes apart from atp8) and recovered 9 individual mitochondrial gene containing contigs from *Dicyema japonicum* and from a second unidentified species (*Dicyema sp.; cox1, 2, 3; cob;* and *nad1, 2, 3, 4, 5*). Cob, nad3, nad4, and nad5 had not previously been identified in any *Dicyma* species. All protostomes studied possess a unique, derived combination of amino acid signatures and conserved deletions in their mitochondrial NAD5 genes. Comparing the NAD5 protein coding regions of *Intoshia* and *Dicyema* to those of other Metazoa shows that both share almost all of the conserved protostome signatures [11] (Fig. 2a). This signature is significantly more complex than the two amino acids of the Lox5/DoxC signature from *Dicyema* previously published [11-13] and shows beyond doubt that both groups are protostomes.

**Fig. 1:**
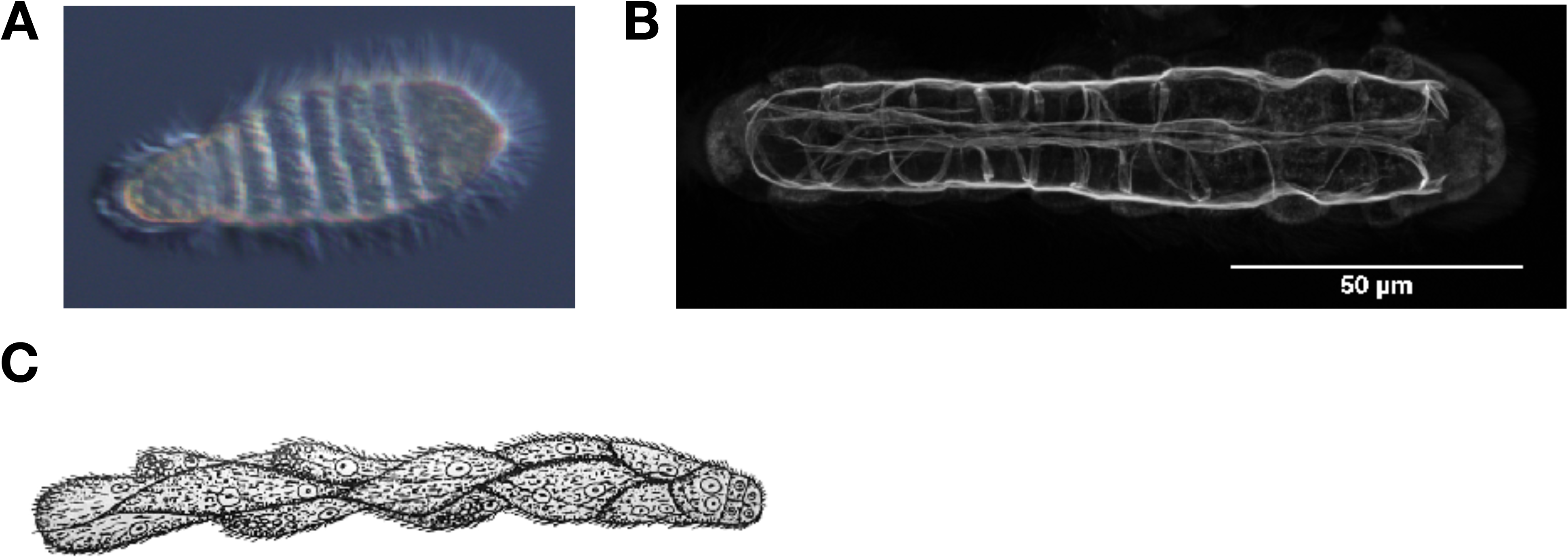
The mesozoans *Intoshia variabili* and *Dicyema typus*. A. Differential Interference contrast micrograph of an *Intoshia variabili* female showing repeated bands of ciliated cells. Picture G. Slyusarev (St Petersburg State University, Russia). B. Confocal image of a phalloidin stained female specimen of *Intoshia linei* reveals repeated set of circular muscles. Picture G. Slyusarev (St Petersburg State Univ.). C. Rhombogen stage of a dicyemid (*Dicyema typus* from the Octopus) adapted from Hyman L.H. The Invertebrates: Protozoa through Ctenophora McGraw-Hill, New York 1940(*19*). Anterior to right in all images.

**Fig. 2:**
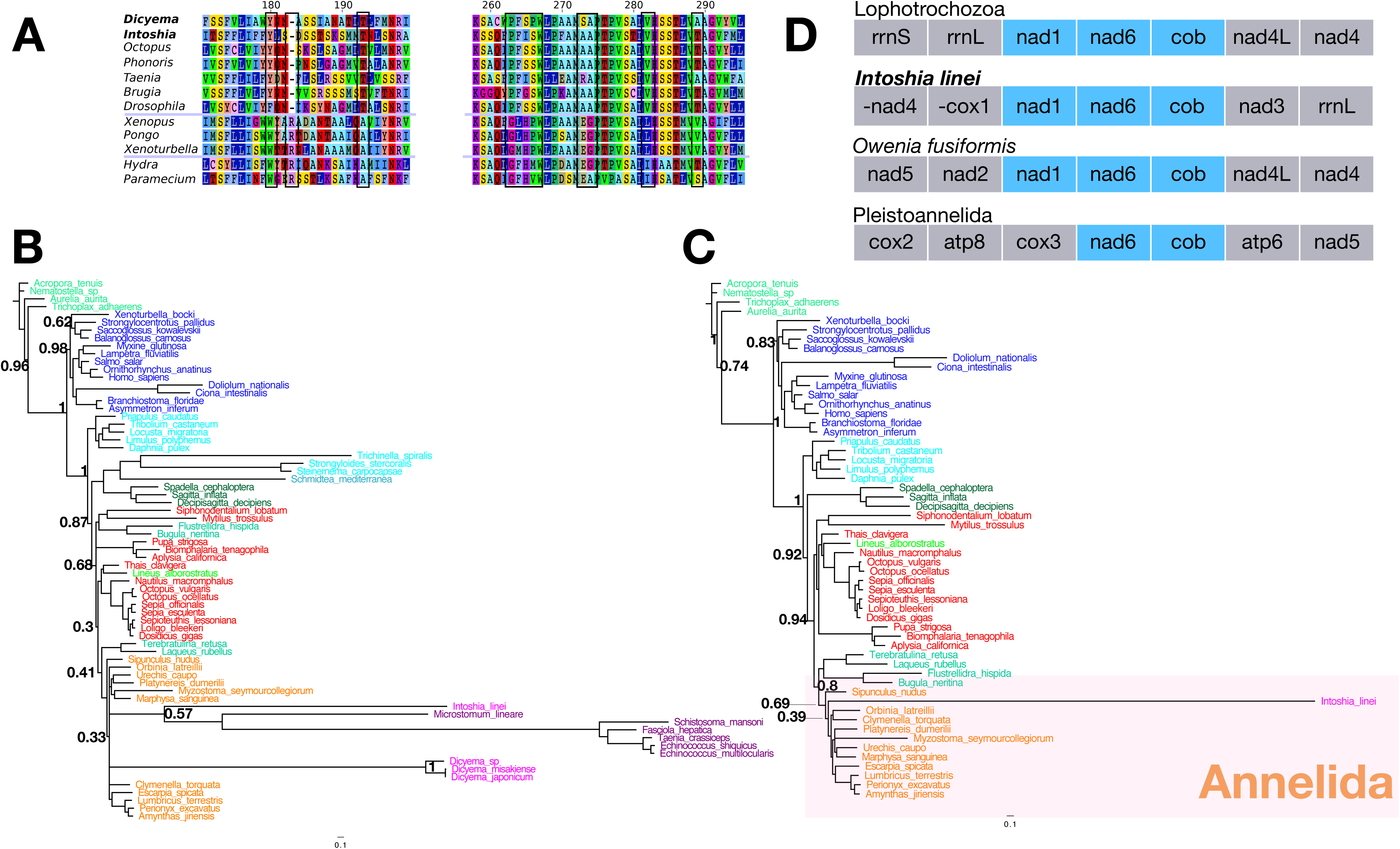
Analyses of the phylogenetic positions of *Dicyema* and *Intoshia* based on mitochondrial gene sequences. A. Alignment of the mitochondrial NAD5 gene from selected protostomes, deuterostomes, and outgroups, highlighting derived substitutions and amino acid deletions shared by the orthonectids, dicyemids, and other protostomes. B. A mitochondrial bayesian phylogeny based on 2969 positions places orthonectids and dicyemids inside Lophotrochozoa, but the unlikely assemblage of *Intoshia linei* and flatworms with annelids suggest this is affected by systematic error. C. Mitochondrial bayesian phylogeny omitting the long branching taxa including *Dicyema* gives some support for a position of *Intoshia* within Annelida.D. Order of the *Intoshia* nad1, nad6, and cob mitochondrial genes in comparison to the early branching annelid *Owenia fusiformis,* the pleistoannelid ground plan and the lophotrochozoan ground plan (see ref [17]).

It has been suggested that mesozoans are derived from the parasitic neodermatan flatworms. If this were correct mesozoans would be expected to share two changes in mitochondrial genetic code that unite all rhabditophoran flatworms, where the triplet AAA codes for Asparagine (N) rather than the normal Lysine (K) and ATA codes for Isoleucine (I) rather than the usual Methionine (M) [14]. We inferred the mitochondrial genetic codes for *Dicyema* and *Intoshia.* Both groups have the standard invertebrate mitochondrial code arguing against a relationship with the parasitic rhabditophoran platyhelminths (table S1).

We next aligned the mitochondrial genes of *Intoshia* and three species of *Dicyema* with orthologs from a diversity of other Metazoans and concatenated these to produce a matrix of 2,969 reliably aligned amino acids from 69 species. Phylogenetic analyses of this comparatively small data set is not expected to be as reliable as a much larger set of nuclear genes and aspects of the topology and observed branch lengths suggest it was affected by LBA (Fig. 2b). To reduce the effects of LBA on the inference of the affinities of the mesozoans we removed the taxa with the longest branches and considered the position of the dicyemids and orthonectid separately (as both are very long branched). We were unable to resolve the position of the dicyemids (although they are clearly lophotrochozoans), but found some support for placing the orthonectid *Intoshia linei* with the annelids (Fig. 2c and figures S1, 2). *Intoshia linei* has a unique mitochondrial gene order although the order of the genes nad1, nad6, and cob match that seen in the Lophotrochozoan ground plan and the early branching annelid *Owenia* (Fig. 2d).

We next assembled a data set of 469 orthologous genes, 227,187 reliably aligned amino acids, from 45 species of animals including *Intoshia linei* and two species of *Dicyema*. After removing positions in the concatenated alignment with less than 50% occupancy we had an alignment length of 190,027 amino acids and average completeness of ~68%. *Intoshia linei* was 65% complete, while *Dicyema japonicum* and *Dicyema sp*. were 77% and 43% complete respectively (table S2). We conducted a bayesian phylogenetic analyses of these data with the site heterogeneous CAT+G4 model in Phylobayes [15]. To provide an additional, conservative estimate of clade support and to enable further analyses in a practical time frame, we also used jackknife subsampling. For each jackknife analysis we took 50 random subsamples of 30,000 amino acids each and ran 2,000 cycles (phylobayes CAT+G4) per sample. All 50 subsamples were summarised into a single tree with the first 1800 trees from each excluded as ‘burnin’ [16].

We observed strong support for a clade of Lophotrochozoa (excluding Rotifers) including both dicyemids and the orthonectid (Bayesian Posterior Probability (PP) = 1.0; Jackknife Proportion (JP) = 0.97) (Fig. 3a). The dicyemids and orthonectids were not each other’s closest relatives; the position of the dicyemids within the Lophotrochozoa was not resolved; they were not the sister group of the platyhelminths nor of the gastrotrichs in our analysis. The position of the orthonectid *Intoshia*, in contrast, was resolved as being within the clade of annelids (Fig. 2a PP = 0.97; JP = 0.74).

**Fig. 3:**
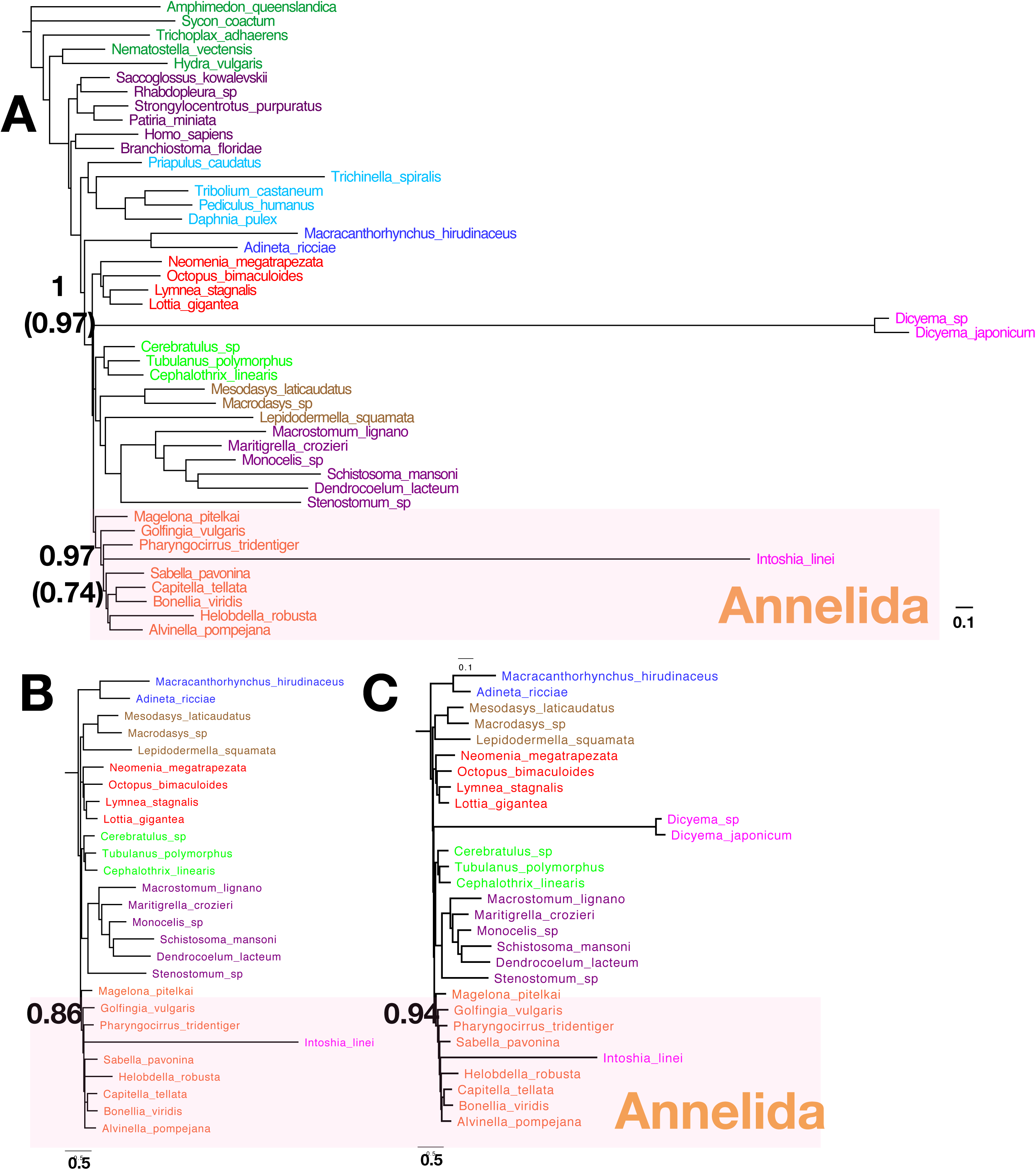
Analyses of the phylogenetic positions of *Dicyema* and *Intoshia* based on nuclear gene sequences. A. A bayesian phylogeny reconstructed from 190,027 aligned amino acid positions analysed under the CAT+G4 model. Support values are from bayesian posterior probabilities (PP) and from 50 jackknifed sub-samples of 30,000 residues (JP support values in brackets). Both analyses reveal Mesozoa to be polyphyletic and place *Intoshia linei* in Annelida (see Supp Fig 4a for support values). B. A repeat of the jackknife analysis omitting the long-branching *Dicyema* species eliminates the potential for LBA between *Intoshia* and *Dicyema*. This leads to an increase in the support for a position of *Intoshia* within Annelida from JP 0.74 to JP 0.86 JP. (Only lophotrochozoan part of the tree shown, see Supplementary Fig 4c for full tree). C. Bayesian jackknife using CAT+G4 model using the best quarter of genes supporting monophyletic annelids leads to increased support for *Intoshia* within Annelida to JP 0.94 even with the inclusion of the *Dicyema* species. (Only lophotrochozoan part of the tree shown, see Supplementary Fig 4d for full tree).

We next asked whether there was any effect from long branched dicyemids on the strength of support for inclusion of *Intoshia* within the Annelida - *Intoshia* also being a long-branched taxon. Repeating our jackknife analyses with dicyemids excluded increased the support for *Intoshia* as an annelid from JP = 0.74 to JP = 0.86 (Fig. 3b) showing that when the expected LBA between *Dicyema* spp and *Intoshia* is prevented, there is stronger support for including the orthonectid in Annelida. An equivalent analysis omitting *Intoshia* did not help to resolve the position of dicyemids (figure S3).

To test further the support for *Intoshia* being a member of Annelida, we reasoned that an analysis restricted to genes showing the strongest signal supporting monophyletic Annelida should give stronger support to *Intoshia* within Annelida but only if it is indeed a member of the clade; if not, support should decrease when using this subset of genes. We first removed all mesozoan sequences from each individual gene alignment and reconstructed a tree for each gene. We ranked these gene trees according to the proportion of all annelids present in a given gene data set that were observed united in a clade. We concatenated the genes (now including mesozoans) from strongest supporters of monophyletic Annelida to weakest. We repeated our jackknife analyses using the best quarter of genes. An analysis of the genes that most strongly support monophyletic Annelida results in an increase support for inclusion of *Intoshia* within Annelida from JP = 0.74 to JP = 0.94 (Fig. 3c).

Our results suggest that recent findings of a close relationship between *Intoshia* and *Dicyema* and the linking of both these taxa to rapidly evolving gastrotrichs and platyhelminths [7,8] is due to long branch attraction. To test this prediction we exaggerated the expected effects of LBA on our own data set by using less well fitting models. We first conducted cross validation comparing the site heterogeneous CAT+G4 model we have used to the site homogenous LG+G4 and show that LG+G4 is a significantly less good fit to our data (CAT+G4 is better than LG+G4: Δln*L* = 9787 +/− 249.265). We used the less well fitting LG+G4 model to reanalyse the jackknife replicates of a data set including our four most complete annelids. We observed a topology clearly influenced by LBA in which long branched taxa including flatworms, annelids, rotifers and nematodes were grouped. We also observed within this ‘LBA assemblage’ the two longest branched clades, dicyemids and the orthonectid as each other’s closest relatives. As a further test we reanalysed the published data set [8] which had linked orthonectid and dicyemid with platyhelminths and gastrotrichs. When we removed the most obvious source of LBA - the long branched dicyemid - we found that the orthonectid *Intoshia* was, as expected, found not with platyhelminths or gastrotrichs but with the two annelids present in this data set, again providing evidence of the effects of long branch attraction (Fig 4).

**Fig. 4:**
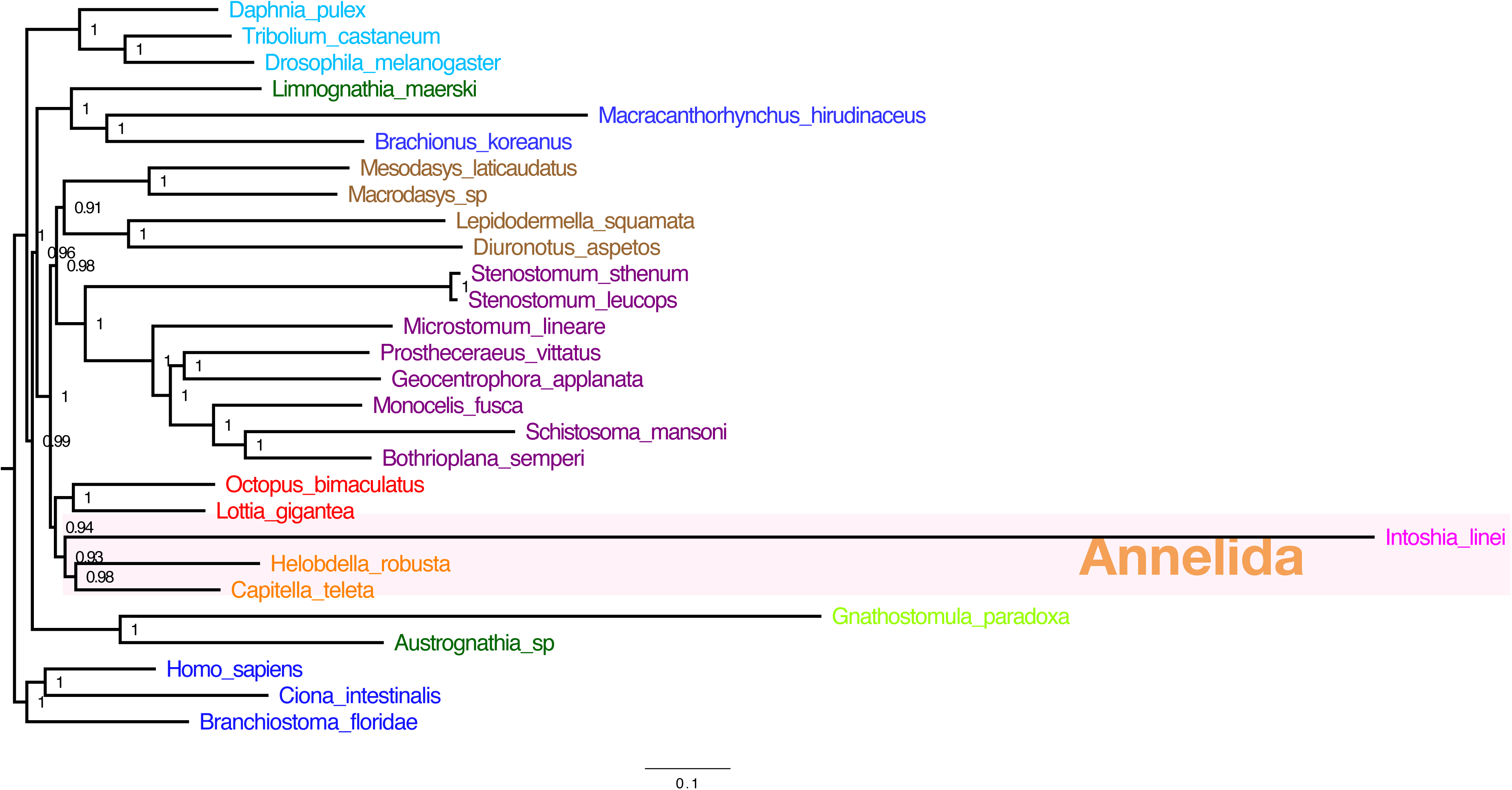
Reanalysis of a published data set addressing potential LBA between mesozoans supports annelid affinity for *Intoshia.* Repeating the analyses on a previously published data set [8] excluding the long branching *Dicyema* leads to *Intoshia* being placed with the annelids, showing the likely effect of LBA on the original analysis. Support values are bayesian posterior probabilities (PP).

## Discussion

We have analysed the first, almost complete mitochondrial genome sequence of an orthonectid mesozoan and added to the known mitochondrial genes of Dicyemida to provide two powerful rare genomic changes. Our analyses of mitochondrial NAD5 gene sequences show unequivocally that both Dicyemida and Orthonectida are members of the protostomes and the absence of rhabditophoran flatworm mitochondrial genetic code changes rejects existing ideas that either group might be derived from parasitic flatworms. Both groups show unusually high rates of evolution and this required steps to test for and avoid the possible effects of long branch attraction, not least between the orthonectids and dicyemids.

Our mitochondrial data set and our large, taxonomically broad set of nuclear genes with a low percentage of missing data, analysed with well fitting, site heterogeneous models of sequence evolution, do not support the close relationship between orthonectids and dicyemids. Orthonectids are annelids and not members of the Mesozoa and the phylum Mesozoa *sensu lato* is an unnatural polyphyletic assemblage. We were unable to place the dicyemids more precisely and they may be considered a phylum in their own right. Experiments manipulating the expected effects of LBA strongly suggest previous phylogenies were affected by this important source of systematic error. Finding the orthonectids and dicyemids not closely associated demonstrates a remarkable instance of convergent evolution in two unrelated, miniaturised parasites.

The finding that the orthonectid *Intoshia* is a member of the Annelida shows that it has evolved its extraordinary simplicity by drastic simplification from a much more complex annelid common ancestor. Our phylogenetic analyses could not more precisely place *Intoshia* within the annelids, however, a short stretch of mitochondrial genes (nad1, nad6, cob) that are found in the same order as in the lophotrochozoan ancestor and in the early branching annelid *Owenia fusiformis* but not in the pleistoannelid ground plan argues for a position outside of the Pleistoannelida [17] (Fig 2d). Possible evidence of an ancestral segmented body plan is still apparent in the series of circular muscles regularly spaced along the antero-posterior axis of *Intoshia* (Fig 1b), along with similarly repeated bands of cilia (Fig 1 and ref [18]). Further analysis of the genome, embryology and morphology of *Intoshia* or other orthonectids are predicted to show additional clues as to their cryptic annelidan ancestry.

## Methods

### Genome and transcriptome assemblies

We downloaded genomic (*Intoshia linei*: SRR4418796, SRR4418797) and transcriptomic (*Dicyema sp*.: SRR827581; *Dicyema japonicum:* DRR057371) data from the NCBI Short Read Archives and DDBJ, and used Trimmomatic [19] to clean residual adapter sequences from the sequencing reads and to remove low quality bases. We used the clc assembly cell (clcBIO/Qiagen; v.5.0) to re-assemble the *I. linei* genome and the Trinity pipeline [20] (v.2.3.2) to assemble the *Dicyema.* sp. and *D. japonicum* transcriptomes using default settings. We additionally assembled transcriptomes for *Phascolopsis gouldii, Spiochaetopterus sp., Arenicola marina, Sabella pavonina, Magelona pitelkai, Pharyngocirrus tridentiger* and *Bonellia viridis* from SRA datasets (SRR1654498, SRR1224605, SRR2005653, SRR2005708, SRR2015609, SRR2016714, SRR2017645) using the same approach.

### Identifying mitochondrial genome fragments

Using mitochondrial gene protein coding sequences from flatworms as queries [21] we used tblastn [22] and blastp to search for *Dicyema* sp. and *D. japonicum* mitochondrial fragments in the Trinity RNA-Seq assemblies, and screened the *I. linei* genome re-assembly in a similar way. Positively identified ORFs were then blasted against NCBI nr to detect possible contamination from host species in the RNA-Seq data. For each *Dicyema* sp. gene-bearing contig, we also found additional contigs which had strongly matching blast hits to *Octopus* or other cephalopods (or in some cases to the gastropod mollusc *Aplysia*) and we discarded these as likely contaminations.

### Annotating mitochondrial genomes

Using blast we identified a 14.2kb mitochondrial contig in the assembled *I. linei* genome, which we annotated using MITOS [23]. The location of protein-coding genes were manually verified from MITOS prediction, and inferred to start from the first in-frame start codon (ATN, GTG, TTG, or GTT). The C-terminal of the protein-coding genes was inferred to be the first in-frame stop codon (TAA, TAG or TGA). We aligned the *Intoshia* and *Dicyema* NAD5 genes with those from 5 protostomes, 4 deuterostomes, and 2 non-bilaterian species in the Geneious software to visualise Protostome specific signatures in the sequence.

### Mitochondrial Phylogenetics

We grouped the mesozoan mitochondrial protein coding genes with their orthologs from 65 other species selected to cover the diversity of the Metazoa including diploblasts, deuterostomes and ecdysozoans but with an emphasis on the diversity of Lophotrochozoa. We aligned each set of orthologs using Muscle [24] v3.8.31 using default parameters and trimmed these alignments to exclude unreliably aligned positions using TrimAl [25] (version 1.2 rev 59 using default settings). Finally, we concatenated the trimmed alignments of all genes into a supermatrix of 2969 positions. We inferred a phylogeny with phylobayes (4.1b) under the CAT+G4 model. We ran 10 independent chains for 10,000 cycles each. We summarised all ten chains (bpcomp) discarding the first 8,000 trees from each as burnin. We reconstructed additional mitochondrial phylogenies omitting (i) the long branching flatworm species, (ii) all long branch taxa and also *Intoshia*, and (iii) long branch taxa and the *Dicyema* species. Here and elsewhere we visualised and edited phylogenetic trees with FigTree (v1.4.3; http://tree.bio.ed.ac.uk/software/figtree/).

### Nuclear gene orthology determination

We chose to add the mesozoan data to sets of orthologous genes that were previously successfully used to infer lophotrochozoan phylogeny [26,27]. We first used Orthofinder [28] (v.1.0.8) to calculate orthologous relationships between the genes predicted for *I. linei* in the recent genome paper [8] and our *Dicyema* sp. gene predictions. To ensure robustness of the analysis we included several outgroup species (Supplementary table 2) In particular, as we were concerned about potential contamination by the hosts of the parasitic *Dicyema* we included the *Octopus bimaculoides* proteome. Since the published phylogenomic studies included few annelid species we added our own Trinity assemblies of several additional species (see above). We then extracted all orthologous groups containing the *Octopus* and the two mesozoan taxa from the Orthofinder output and inserted these sequences into the original alignments. This resulted in 590 orthologous groups. With the aid of OMA [29] and custom Perl scripts we filtered these groups to contain single copy orthologs of all species. We realigned each set of orthologs using clustal-omega [30]; we removed unreliably aligned positions from each alignment using TrimAl; finally we constructed individual gene trees from these trimmed alignments using phyml [31] (v20160207). Using Python code and the ETE3 toolkit we checked each tree for instances where sequences from *Octopus* and *Dicyema* sp. were each other’s closest relatives (suggesting the sequence is an *Octopus* contaminant) and removed the 5 alignments where the trees had this topology from our set. We concatenated all single trimmed alignments of 45 taxa into a supermatrix of 227,646 positions. We used a custom script to eliminate all positions in the alignment with less than 50% occupancy.

### Nuclear Gene Phylogenomic analyses

Using the mpi version of phylobayes (in v.1.7) run over four independent chains for 5000 cycles and discarding the first 4500 trees as burnin we reconstructed a phylogeny using this alignment under both the CAT+G4 model of molecular evolution. To provide a conservative measure of clade support and to test different data samples in a reasonable time we also reconstructed trees using 50 jackknife sub-samples of 30,000 positions each from the supermatrix. We used phylobayes 4.1c with the aid of the gnu-parallel command line tool [32] and the UCL HPC cluster. We used the CAT+G4 model, and also compared results from LG+G4. We ran phylobayes for 2000 cycles per jackknife sample which consistently resulted in a plateauing of the likelihood score. We summarised all 50 of these phylobayes analyses per model (using bpcomp) discarding the first 1800 sampled trees per jackknife as burnin. We also tested the effect of different species compositions in our dataset by performing phylobayes jackknife sampling with different subsets of taxa.

### Cross validation

We compared the fit of CAT+G4 and LG+G4 models to our data using cross validation as described in the phylobayes user manual. We ran 10 replicates and for each replicate we used a randomly selected 30,000 positions of the data as a training set and 10,000 randomly selected positions as the test set. Log likelihood scores were averaged over the ten replicates using the sumcv command.

### Ranking genes according to support for monophyletic Annelida

We first removed all *Intoshia* and *Dicyema* sequences from each individual gene alignment. For each individual gene, we reconstructed a tree from the aligned protein coding sequences using Ninja [33]. Each tree was parsed using a custom script to find the proportion of annelids in the data set present in the largest clade of annelids found. The tree was given a score which was calculated as the number of annelids in the largest clade/total number of annelids on the tree. Trees with larger monophyletic annelid clades scored highest. The genes were then concatenated in order of their score. We took the first 25% of positions from this concatenation (those genes with the strongest signal supporting monophyletic annelids) and analysed jackknife replicates as before.

## Acknowledgements

The authors are grateful to Prof George Slyusarev (Saint Petersburg State University, Russia) for providing images of *Intoshia linei* and for sharing an unpublished book chapter and to Dr Tsai-Ming Lu and colleagues (Okinawa Institute of Science and Technology, Japan) for sharing data. The authors also thank Fraser Simpson (UCL) for help in editing Figure 1b. The research was funded by ERC grant (ERC-2012-AdG 322790) to MJT. Alignments have been deposited with Zenodo (XXX) and phylogenetic trees are available through treebase.org (XXX).

## Author Contributions

Conceived the study: MJT. Planned the study: MJT and PHS. Assembled the data sets: PHS. Analysed the data: PHS and MJT. Drafted the manuscript: MJT and PHS. Analysed mitochondrial data: HER.

## Supplementary Tables

**Supplementary Table 1**

Predicted correspondence of nucleotide triplets to amino acids in *Intoshia* and three *Dicyema* species. For each triplet, the amino acid corresponding to the triplet in the standard invertebrate mitochondrial code is shown, the number of observations of the triplet to prediction is based on, the predicted amino acid and its score and finally the second highest scoring amino acid prediction. The triplets AAA and ATA are highlighted in green and likely errors highlighted in blue. Likely errors are mostly associated with very low numbers of observed GC rich triplets in these very AT rich mitochondrial genomes.

**Supplementary Table 2**

List of species used in the final phylogenetic analysis, data sources, and representation in the final alignment.

## Supplementary Figures

**Figure S1. Related to Figure 2.**
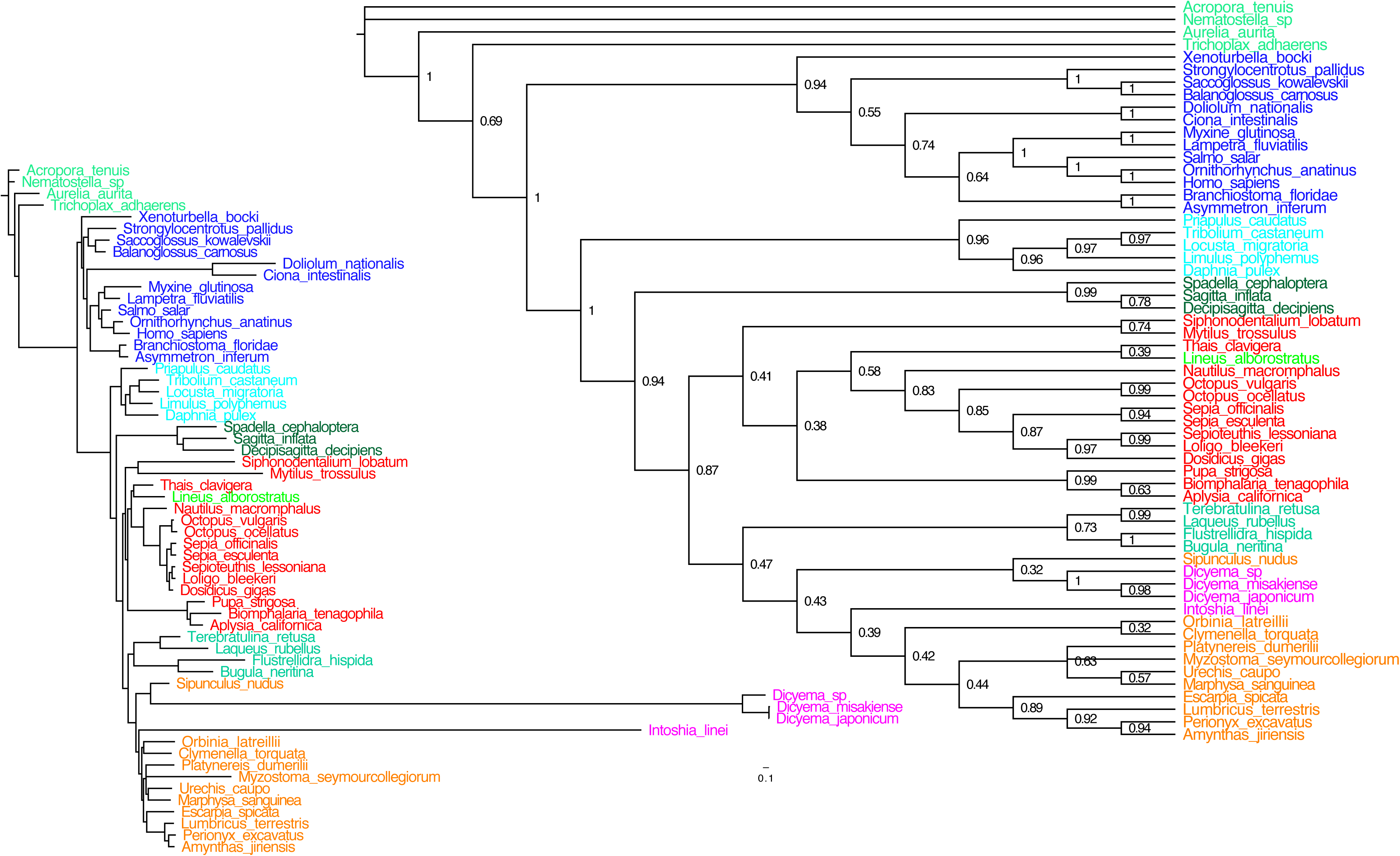
Phylogram and corresponding cladogram of a Bayesian analysis of our mitochondrial data set omitting the long-branching flatworm species. Phylobayes CAT+G4 model was run in 10 independent runs for 10,000 cycles each on an alignment with 2969 positions and 8000 trees were discarded as burnin.

**Figure S2. Related to Figure 2.**
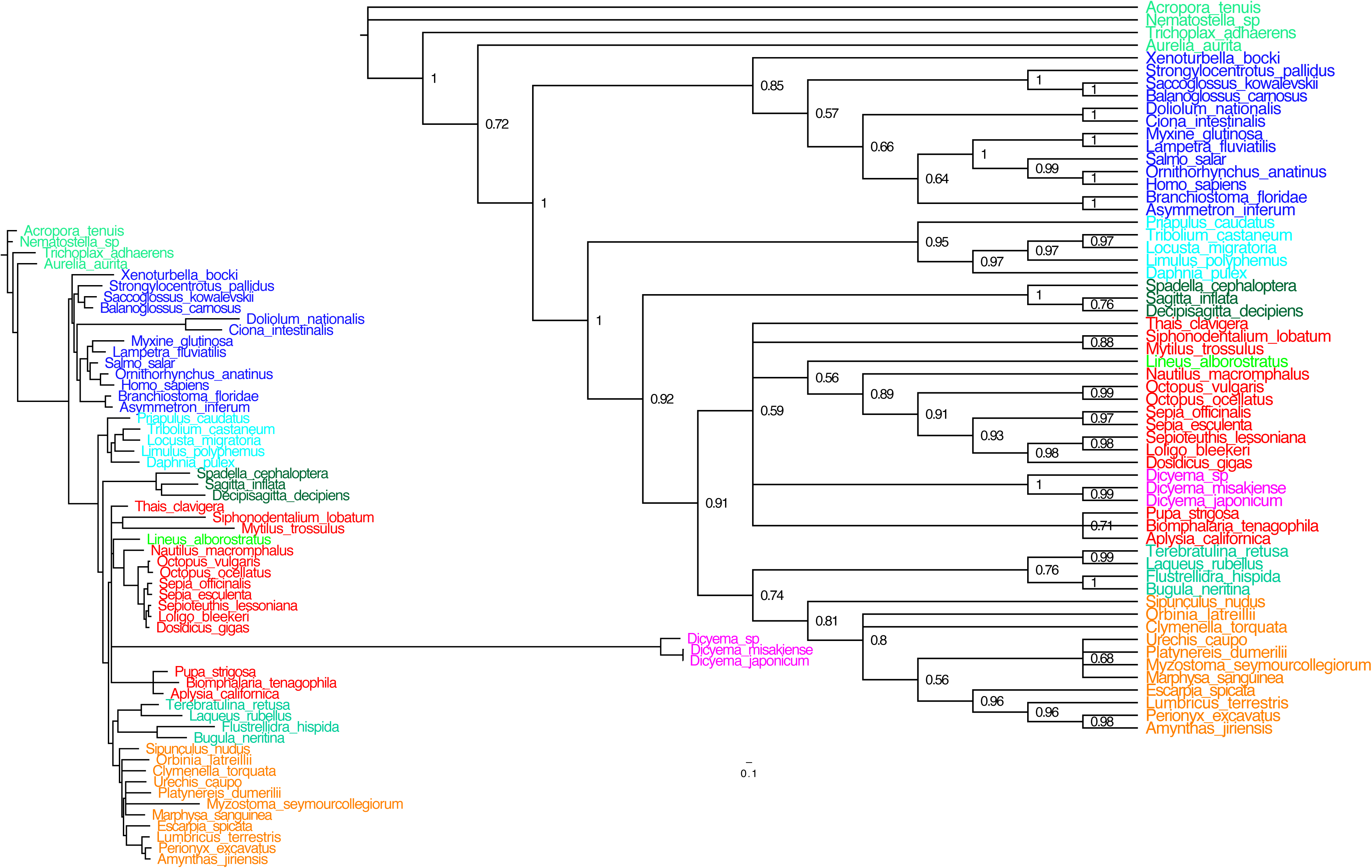
Phylogram and corresponding cladogram of a Bayesian analysis of our mitochondrial data set omitting *Intoshia linei.* Phylobayes CAT+G4 model was run in 10 independent runs for 10,000 cycles each on an alignment with 2969 positions and 8000 trees were discarded as burnin.

**Figure S3. Related to Figure 3.**
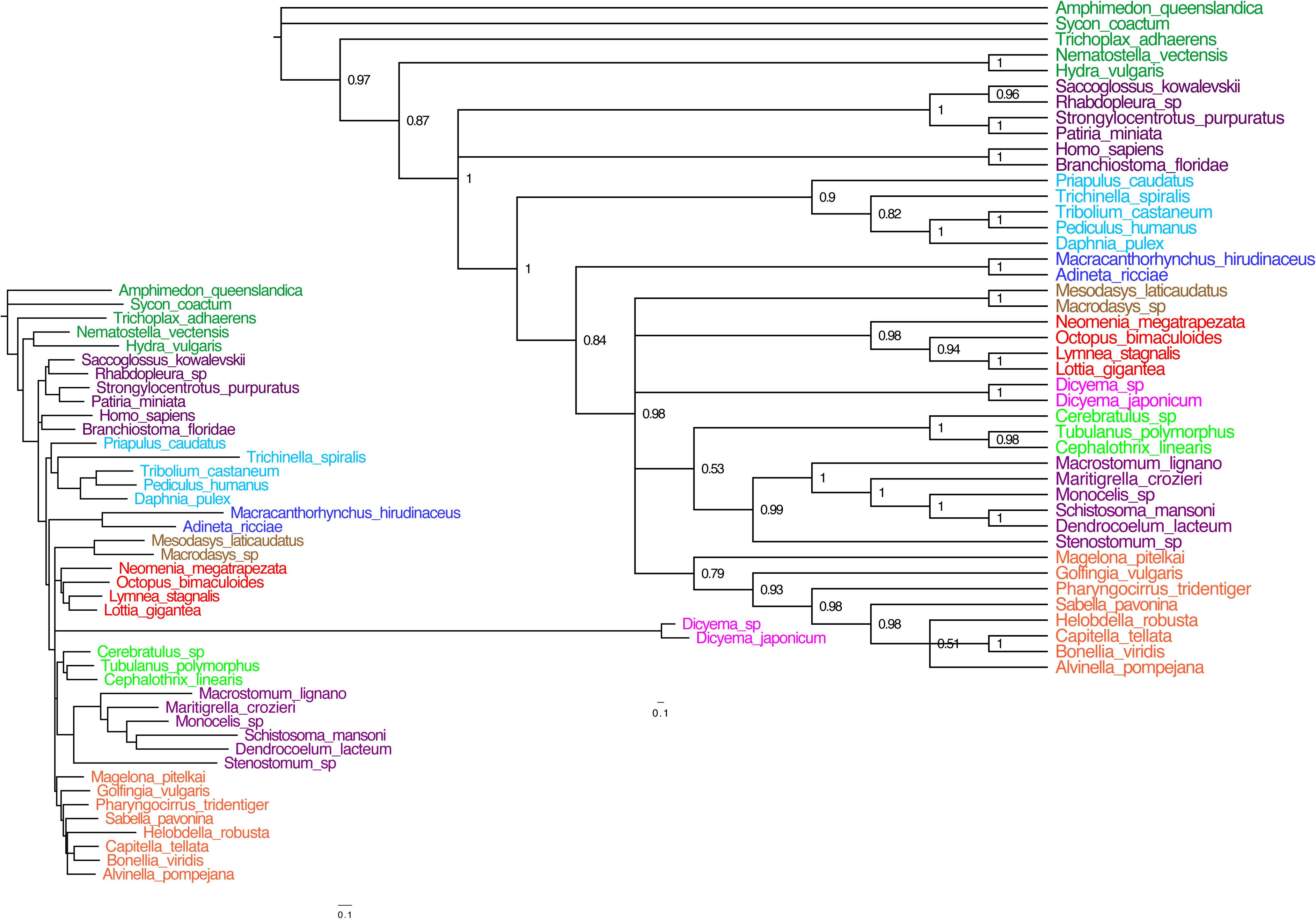
A phylogram based on our analysis of the jackknifed dataset omitting *Intoshia linei*. Contrary to the improvement in placing *I. linei* observed when excluding the *Dicyema* species, the exclusion of *I. linei* does not lead to a better resolution of the *Dicyema* species’ position. This can be seen as further evidence for the non-affiliation of orthonectids and dicyemids and the correct inference that orthonectids are part of Annelida.

**Figure S4. Related to Figure 3.**
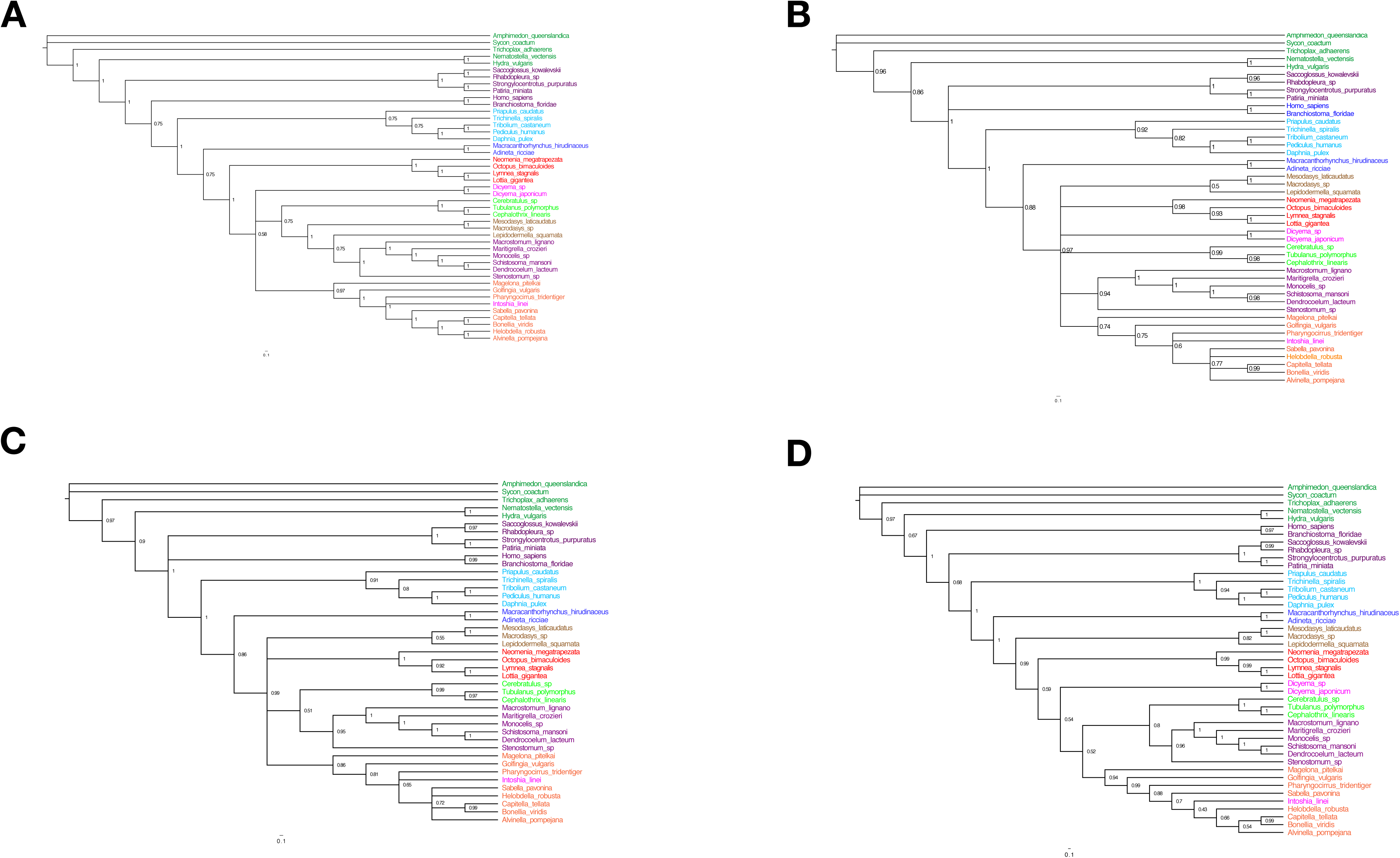
A. Cladogram corresponding to Fig 3a showing all PP support values for the CAT+G4 phylogeny based on the full alignment of 190,027 amino acid positions. B. A cladogram including JP support values based on 50 jackknife subsamples of 30,000 amino acid positions each independently analysed for 2000 cycles under the CAT+G4 model in phylobayes and summarised with the bpcomp command setting 1800 as burnin. As in the analysis of the full dataset I. linei is found within the annelids and phylum Mesozoa is found as an unnatural assemblage. C. Cladogram corresponding to Fig 3b showing all support values. D. Cladogram corresponding to Fig 3c showing all support values.

